# A luminescent attenuated SARS-CoV-2 for the identification and validation of drug-resistant mutants

**DOI:** 10.1101/2025.05.09.653029

**Authors:** Yao Ma, Chengjin Ye, Ahmed Magdy Khalil, Sara H. Mahmoud, Elizabeth B. Sobolik, Alexander L Greninger, Esteban Castro, Nathaniel Jackson, Mahmoud Bayoumi, Richard K Plemper, Luis Martinez-Sobrido

## Abstract

The emergence of SARS-CoV-2 variants has necessitated continuous updating of vaccines. In contrast, antivirals remained effective as they target conserved viral proteins that are essential for the viral life cycle. However, several mutations in SARS-CoV-2 that may affect the efficacy of United States (US) Food and Drug Administration (FDA)-approved antivirals have been recently identified. Detecting drug-resistant SARS-CoV-2 mutants and investigating their escape mechanism(s) are critical to guide the selection of effective antiviral therapies. In this study, we constructed an attenuated recombinant (r)SARS-CoV-2 lacking the open reading frame (ORF) proteins 3a and 7b but expressing nanoluciferase (Nluc), rSARS-CoV-2 Δ3a7b-Nluc, to facilitate tracking viral infection. Using this virus, we selected drug-resistant mutants to the main viral protease (Mpro) inhibitor nirmatrelvir. After passaging Δ3a7b-Nluc 10 times in the presence of increasing concentrations of nirmatrelvir, a virus population with enhanced resistance was selected. We identified two non-synonymous mutations (L50F and R188G) in Mpro, encoded by the non-structural protein 5 (NSP5) gene. Using reverse genetics, we generated rSARS-CoV-2 Δ3a7b-Nluc containing the identified L50F and R188G mutations, individually or in combination, and assessed their contribution to nirmatrelvir resistance. Our results indicate that both mutations are involved in escaping from nirmatrelvir. Altogether, our results demonstrate the feasibility of using rSARS-CoV-2 Δ3a7b-Nluc variant to identify and validate mutations that confer resistance to FDA-approved antiviral drugs without the concern of conducting gain of function (GoF) experiments with wild-type (WT) forms of SARS-CoV-2.

**IMPORTANCE:** Small-molecule antiviral drugs have been used for the treatment of SARS-CoV-2 infections. However, drug-resistant SARS-CoV-2 mutants to currently US FDA-approved Mpro targeting antivirals have been identified. Information on SARS-CoV-2 escape mutants and mutations affecting the antiviral activity of licensed antivirals remain limited. In this study, we developed a nanoluciferase (Nluc)-expressing attenuated recombinant (r)SARS-CoV-2 lacking the ORF 3a and 7b proteins (Δ3a7b-Nluc) to identify nirmatrelvir resistant mutants without the biosafety concerns associated with gain-of-function (GoF) research using wild-type (WT) SARS-CoV-2. Using Δ3a7b-Nluc, we have selected variants with reduced sensitivity to nirmatrelvir that were validated by the generation of rSARS-CoV-2 Δ3a7b-Nluc containing the candidate L50F and R188G mutations in Mpro. These results demonstrate the feasibility of using rSARS-CoV-2 Δ3a7b-Nluc to safely identify and validate drug-resistant mutants overcoming concerns originating from adaptation studies using WT SARS-CoV-2.

## INTRODUCTION

Over the past five years, severe acute respiratory syndrome coronavirus 2 (SARS-CoV-2) has caused around 7 million deaths (1). While the burden of coronavirus disease 2019 (COVID-19) has been substantial, vaccines implemented early in 2021 provided some level of protection against viral infection, yet SARS-CoV-2 is still evolving (2–5) and vaccines developed against the original 2019 SARS-CoV-2 strain no longer protect against newly emerging viral variants. Vaccines have been updated to combat the challenge of the latest circulating SARS-CoV-2 strains (6–10). Similarly, monoclonal antibodies with Emergency Use Authorization (EUA) in the United States (US) became rapidly ineffective against SARS-CoV-2 variants and the authorization for clinical use was withdrawn in early 2023 (11–13). Resistance caused by mutations in conserved viral proteins has also been reported for small-molecule antivirals such as nirmatrelvir, ensitrelvir, and remdesivir, though each retains efficacy against the vast majority of circulating strains and no resistance to date has been described for molnupiravir (14–19). To guide the selection of new antiviral therapies and assess their effectiveness against new potentially emerging strains, isolating drug-resistant mutants (DRMs) is a critical first step, which is followed by identifying the mutations that contribute to resistance.

The 3C-like protease (3CLpro), also known as the main protease (Mpro), encoded by the viral nonstructural protein 5 (NSP5) gene, is a cysteine protease responsible for processing SARS-CoV-2 polyproteins open reading frames (ORF) 1a and 1b into 12 functional proteins through cleavage at 11 conserved sites (20, 21). As a critical enzyme for viral replication with high conservation among β-coronaviruses, Mpro represents a promising target for the development of antiviral therapeutics (22). Paxlovid, composed of nirmatrelvir and ritonavir, is the first orally available Mpro inhibitor that was approved for treatment of SARS-CoV-2 infection (23). The major antiviral compound, Nirmatrelvir was originally identified as a covalent peptidomimetic inhibitor of SARS-CoV Mpro during the 2002-2004 outbreak (24–27). Several mutations leading to nirmatrelvir resistance have been identified in NSP5 gene (28–31). For instance, L50F+E166V mutations have led to a 12-to-80-fold decrease in nirmatrelvir effectiveness and an E166V substitution alone has led to a 25-to-288-fold lower potency (28, 29). Importantly, these mutations were also found in immunocompromised patients that received prolonged paxlovid treatment and experienced antiviral failure (32, 33). These findings emphasize the importance of identifying DRMs for guiding the selection of effective antiviral therapies.

Although the isolation of DRMs using wild-type (WT) SARS-CoV-2 can provide critical insights into mutations associated to drug resistance, experiments for the selection of DRMs against US Food and Drug Administration (FDA)-authorized antivirals using WT SARS-CoV-2 are considered gain of function (GoF) research. Thus, an approach to extract critical insight into the viral escape landscape without biosafety concerns is urgently needed. For this purpose, an attenuated form of SARS-CoV-2 emerged as a safe alternative (34). Previous studies from our laboratory have demonstrated that a recombinant (r)SARS-CoV-2 lacking two accessory ORF proteins, 3a and 7b, rSARS-CoV-2 Δ3a7b, was attenuated in K18-hACE2 transgenic mice and golden Syrian hamsters (34).

In this study, we generated a rSARS-CoV-2 Δ3a7b-based reporter virus expressing nanoluciferase (Δ3a7b-Nluc) that we used to identify nirmatrelvir-resistant mutants (DRM-N) by serially passaging the attenuated virus in the presence of increasing drug concentrations. Next-generation sequencing (NGS) analysis of DRM-N identified two mutations, L50F and R188G, within NSP5 gene. To validate the role of these two amino acid changes in conferring resistance to nirmatrelvir, we generated rSARS-CoV-2 Δ3a7b-Nluc containing L50F and R188G, alone or in combination. Our findings demonstrate the contribution of L50F and R188G mutations to nirmatrelvir resistance and validate the use of rSARS-CoV-2 Δ3a7b-Nluc for resistance-profiling of antivirals against SARS-CoV-2, alleviating biosafety concerns of performing these experiments with WT SARS-CoV-2.

## RESULTS

### Generation and characterization of rSARS-CoV-2 Δ3a7b-Nluc

Previous studies from our laboratory demonstrated that a rSARS-CoV-2 lacking the ORF 3a and 7b proteins was attenuated in K18-hACE2 mice and golden Syrian hamsters (34). To generate a luminescent version of this attenuated virus, a nanoluciferase (Nluc) reporter gene was inserted upstream of the viral nucleocapsid (N) gene into a bacterial artificial chromosome (BAC) containing a cDNA copy of the rSARS-CoV-2 Δ3a7b genome (**Fig. 1A**). The resulting rSARS-CoV-2 Δ3a7b-Nluc (referred to as Δ3a7b-Nluc) was rescued in Vero cells expressing hACE2 and TMPRSS2 (Vero-AT) according to our previously described protocol (35–37). The plaque morphology of Δ3a7b-Nluc was evaluated and compared to Δ3a7b in Vero-AT cells. Both live-attenuated viruses (Δ3a7b and Δ3a7b-Nluc) exhibited smaller plaque size than rSARS-CoV-2 WT (**Fig. 1B**), but addition of Nluc substrate made Δ3a7b-Nluc plaques visible (**Fig. 1B**). The growth kinetics of Δ3a7b-Nluc was compared to Δ3a7b and rSARS-CoV-2 WT in Vero-AT cells infected at a multiplicity of infection (MOI) of 0.01 plaque forming units (PFU)/cell. Viral replication of Δ3a7b-Nluc was comparable to that of Δ3a7b, and both recombinants showed reduced replication compared to rSARS-CoV-2 WT (**Fig. 1C**), consistent with our previous observations (34). As expected, Nluc activity was present only in cell culture supernatants from Vero-AT cells infected with Δ3a7b-Nluc and peaked 48 hours post-infection in alignment with viral titers (**Fig. 1D**). These results confirm feasibility of generating a Nluc-expressing Δ3a7b that is replication competent, which could also be used as screening strain in antiviral drug discovery campaigns. We next evaluated the half maximal effective concentration (EC_50_) of nirmatrelvir and remdesivir in Vero-AT cells in comparison with Δ3a7b. Both Δ3a7b-Nluc and Δ3a7b returned similar EC_50_ values for nirmatrelvir (**Fig. 2A**) and remdesivir (**Fig. 2B**), which were comparable to those of SARS-CoV-2 WT described in the literature (38). These results suggest that Δ3a7b-Nluc could be used to identify antivirals with a similar EC_50_ as those observed with SARS-CoV-2 WT.

**Figure 1.**
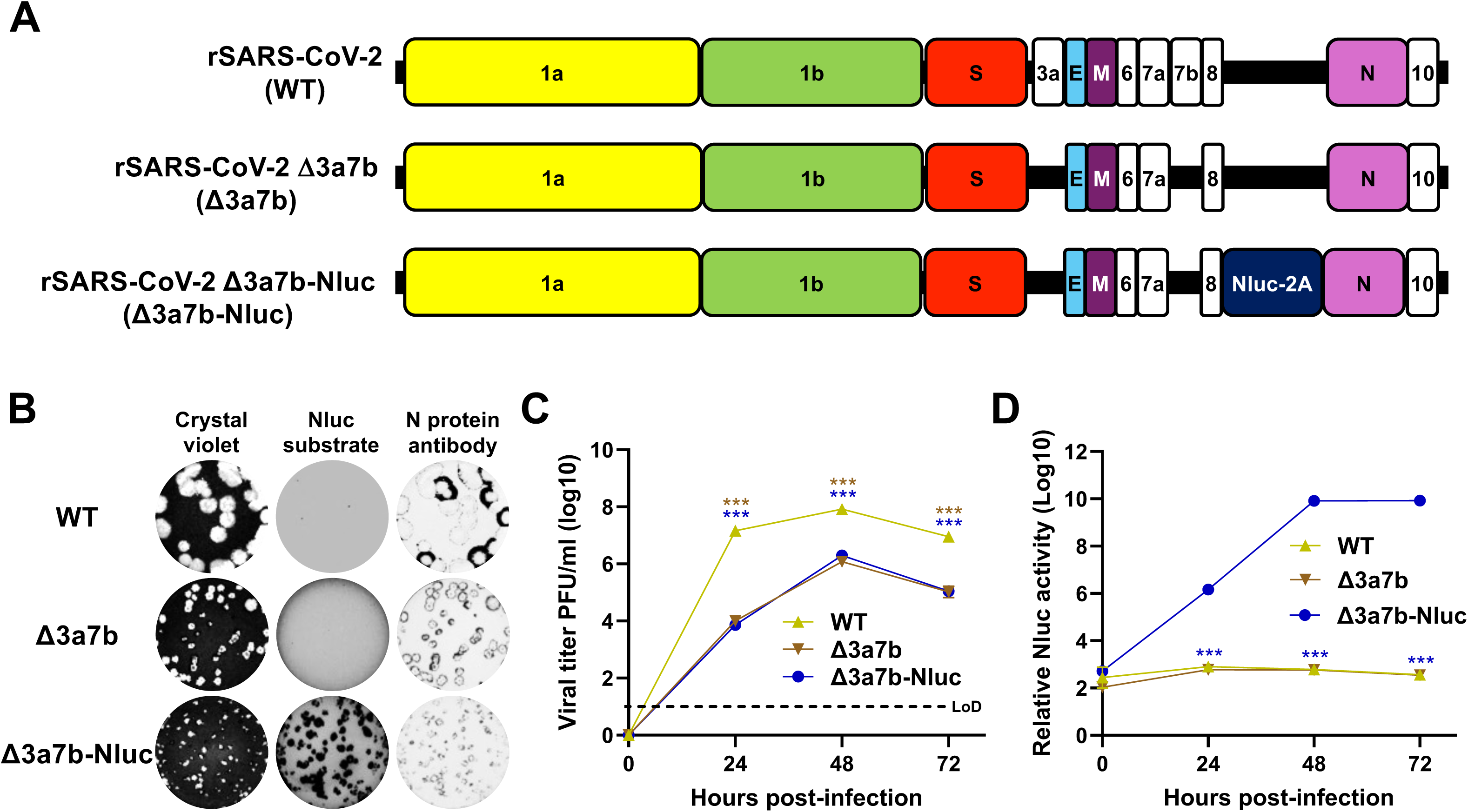
Generation and *in vitro* characterization of Δ3a7b-Nluc. (A) Schematic representation of the viral genomes of rSARS-CoV-2 WT (top), Δ3a7b (middle), and Δ3a7b-Nluc (bottom). (B) The plaque phenotype of rSARS-CoV-2 WT (top), Δ3a7b (middle), and Δ3a7b-Nluc (bottom) in Vero-AT was determined by crystal violet staining (left), Nluc expresssion (middle), and N protein staining (right). (C) Vero-AT cells (6-well plate format, triplicates) were infected (MOI 0.01) with rSARS-CoV-2 WT (yellow), Δ3a7b (brown), or Δ3a7b-Nluc (blue). At the indicated hours post-infection, cell culture supernatants were collected, and presence of virus was determined by plaque assay. LoD: limit of detection. (D) Cell culture supernatants from (C) were used to evaluate the presence of Nluc. Data are presented as mean ± SD. The statistical significance was analyzed with one-way ANOVA followed by Tukey’s test (*** P < 0.001).

**Figure 2.**
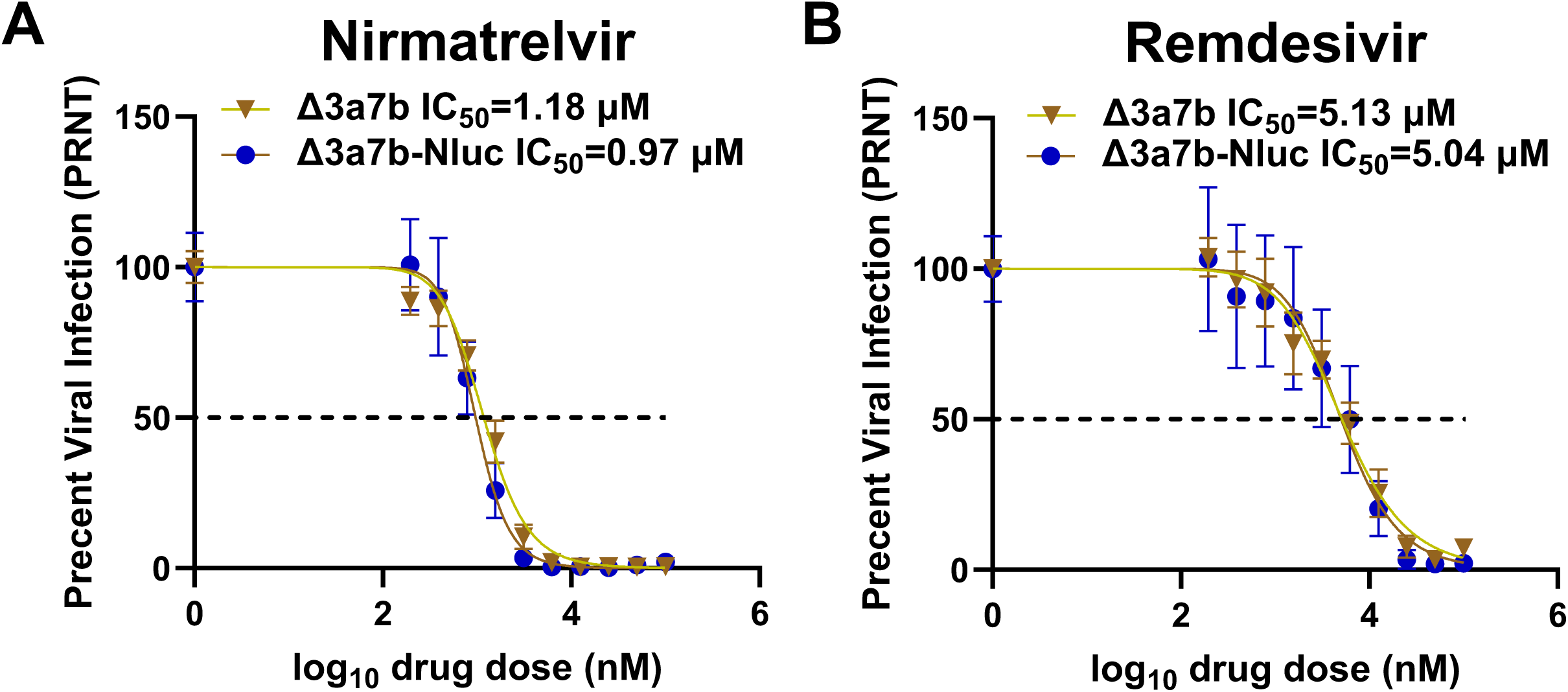
Antiviral activity of nirmatrelvir and remdesivir against Δ3a7b-Nluc. The antiviral activity of nirmatrelvir (A) and remdesivir (B) against Δ3a7b and Δ3a7b-Nluc in Vero-AT cells (96-well plate format, quadruplicates) was evaluated using a PRNT assay. EC_50_ values of the different antivirals were calculated using GraphPad Prism. Data are presented as means ± SD. Dotted line indicates the 50% of viral inhibition.

### Selection of nirmatrelvir drug resistant mutant (DRM-N)

To select viruses resistant to nirmatrelvir, Δ3a7b-Nluc was serially passaged in Vero-AT cells under increasing concentrations of nirmatrelvir (**Fig. 3A**). After 10 serial passages, a virus with enhanced drug resistance to nirmatrelvir (DRM-N) was isolated. The resistance to nirmatrelvir was verified in Vero-AT cells infected with P0 or P10 Δ3a7b-Nluc using an immunofluorescence assay (**Fig. 3B**). Nirmatrelvir at 2.5 µM failed to inhibit P10 Δ3a7b-Nluc DRM-N replication yet resulted in partial inhibition of P0 Δ3a7b-Nluc (**Fig. 3B**). Increased concentrations of nirmatrelvir (10 µM and 40 µM) resulted in complete inhibition of P0 Δ3a7b-Nluc and only partial inhibition of P10 Δ3a7b-Nluc DRM-N replication (**Fig. 3B**). Notably, both Δ3a7b-Nluc P0 and P10 replicate to comparable levels in the absence of drug (**Fig. 3B**). To quantify the resistance of P10 Δ3a7b-Nluc DRM-N, we followed a three-pronged approach: determine plaque reduction in neutralization tests (PRNT), monitor crystal violet staining of infected cells (OD_560_), and quantify Nluc activity (Nluc). PRNT revealed an approximately 14-fold increase in nirmatrelvir resistance for P10 Δ3a7b-Nluc DRM-N (EC_50_ = 12.88 µM compared to P0 Δ3a7b-Nluc EC_50_ = 0.95 µM) (**Fig. 4A**). The OD_560_ results revealed ∼23-fold increase in nirmatrelvir resistance for P10 Δ3a7b-Nluc DRM-N (EC_50_ = 13.96 µM compared to P0 Δ3a7b-Nluc EC_50_ = 0.61 µM) (**Fig. 4B**). The Nluc assay showed a comparable ∼15-fold increase in nirmatrelvir resistance for P10 Δ3a7b-Nluc DRM-N (EC_50_ = 9.18 µM compared to P0 Δ3a7b-Nluc EC_50_ = 0.63 µM) (**Fig. 4C**). As a control, resistance of P0 and P10 Δ3a7b-Nluc DRM-N to remdesivir was evaluated. P10 Δ3a7b-Nluc DRM-N exhibited comparable resistance to remdesivir as P0 Δ3a7b-Nluc across the three assays (**Figs. 4D-4F**), indicating that the acquired resistance was specific to nirmatrelvir. These results suggest that serial passaging of Δ3a7b-Nluc under increasing concentrations of nirmatrelvir resulted in enhanced resistance to the antiviral while maintaining comparable sensitivity to remdesivir, a polymerase inhibitor. Notably, the Nluc-based assay provided consistent EC_50_ values that are comparable to traditional PRNT or OD_560_ assays, while offering a more efficient approach to identify drug resistant mutants.

**Figure 3.**
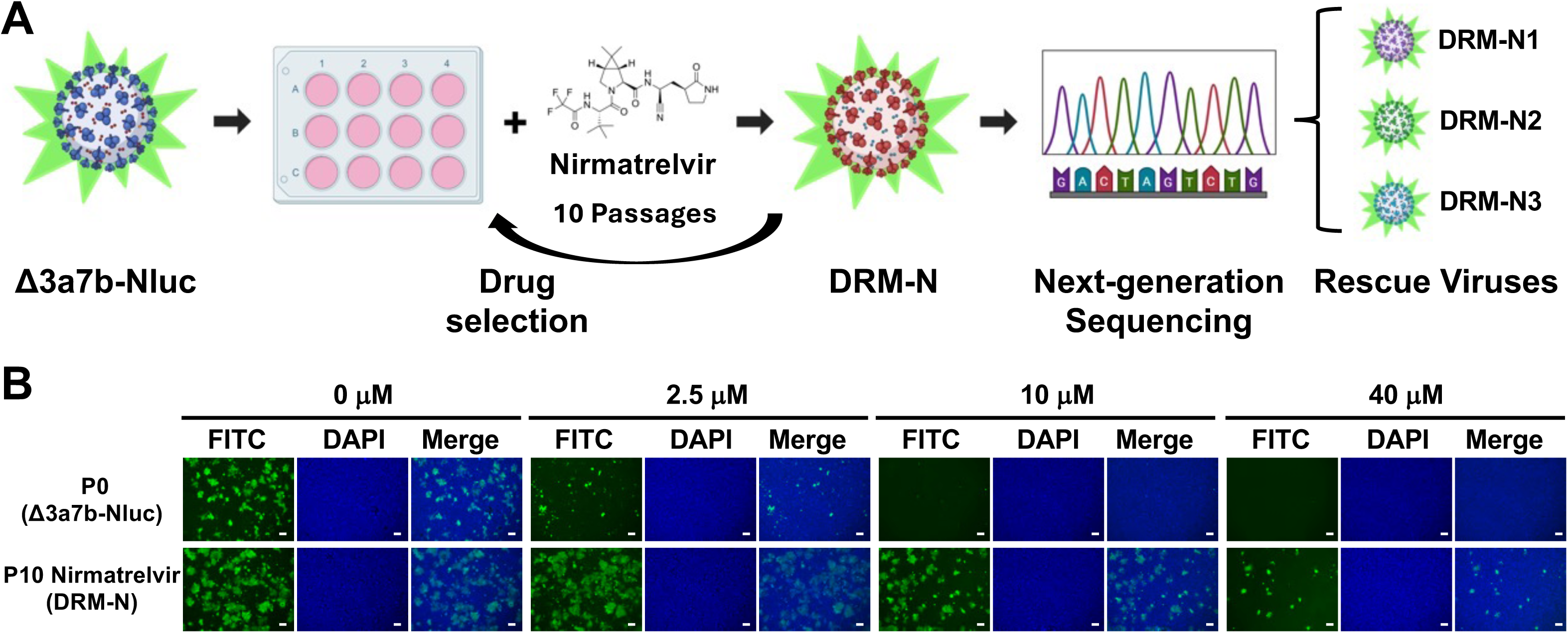
Selection of nirmatrelvir drug-resistant mutants (DRM-N). (A) 3a7b-Nluc was cultured in the presence of increasing concentrations of nirmatrelvir for 10 serial passages (P0-P10). RNA from Δ3a7b-Nluc P10 DRM-N was isolated and used for NGS and Sanger sequencing to identify DRM-N. Then, Δ3a7b-Nluc containing the identified individual and combined mutations were rescued using BAC-based reverse genetics. (B) Vero-AT cells (6-well plate format) were infected (MOI 0.001) with P0 Δ3a7b-Nluc (top) and P10 Δ3a7b-Nluc DRM-N (bottom). After viral adsorption, infectious media was replaced by post-infection media containing the indicated concentrations of nirmatrelvir (0 µM, 2.5 µM, 10 µM and 40 µM). At 48 hours post-infection, cells were fixed, permeabilized, and the presence of viral N protein was determined using the SARS-CoV 1C7C7 cross-reactive monoclonal antibody and an anti-mouse FITC-conjugated secondary antibody. The cell nucleus was stained with DAPI. Scale bars = 200µm.

**Figure 4.**
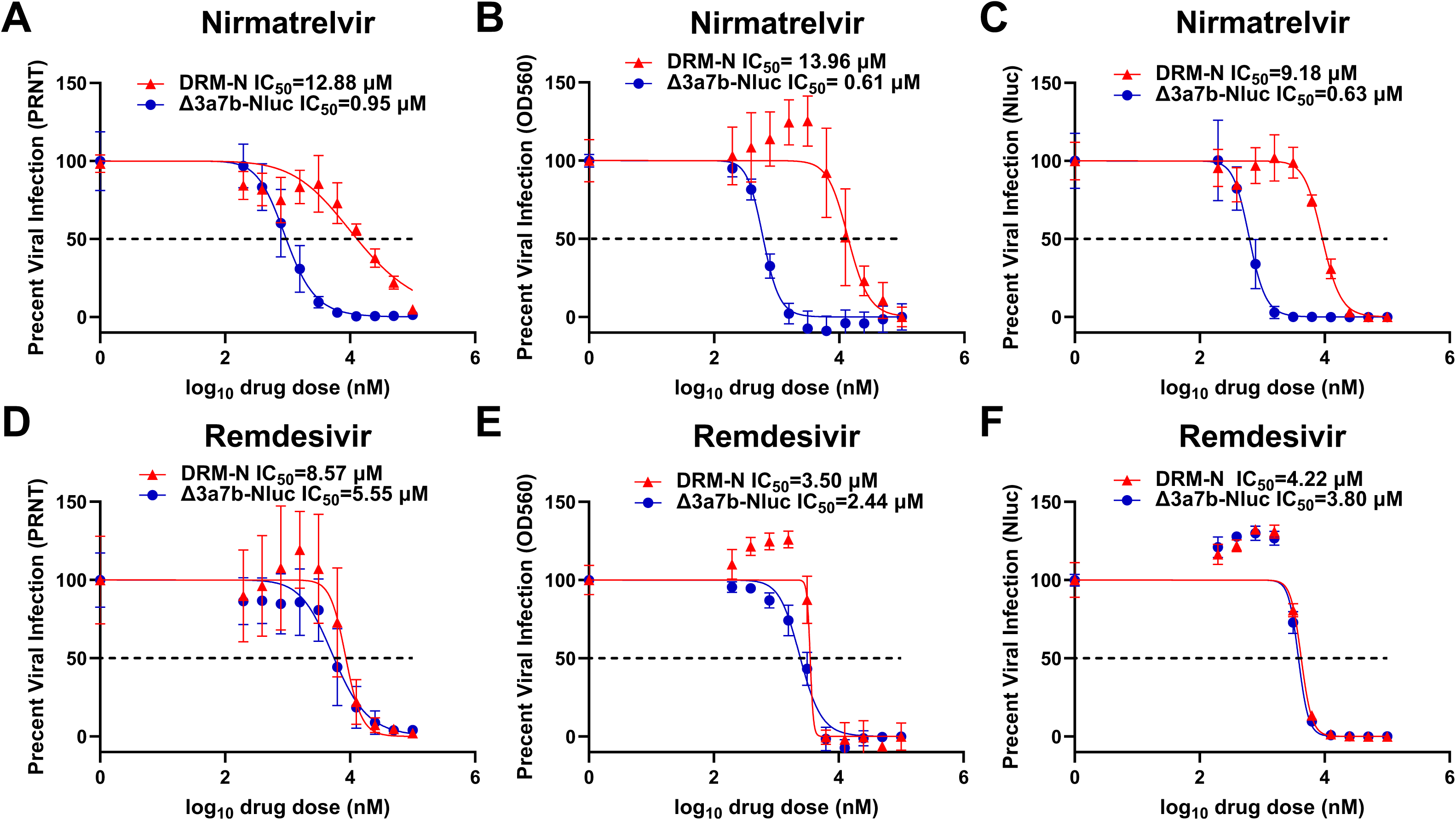
P10 Δ3a7b-Nluc DRM-N resistance to nirmatrelvir and remdesivir. The antiviral activity of nirmatrelvir (A-C) and remdesivir (D-E) against parental (blue) and P10 DRM-N (red) Δ3a7b-Nluc in Vero-AT cells (96-well plate format, quadruplicates) was evaluated using PRNT (A and D), OD_560_ (B and E), and Nluc (C and F) assays. EC_50_ values of the different antivirals were calculated by using GraphPad Prism. Data are presented as means ± SD. Dotted line indicates the 50% of viral inhibition.

### Identification of the amino acids responsible for P10 Δ3a7b-Nluc DRM-N

To identify the mutations responsible for resistance to nirmatrelvir in P10 Δ3a7b-Nluc DRM-N, RNA samples from Vero-AT cells infected with P10 Δ3a7b-Nluc DRM-N were analyzed by NGS (**Fig. 5A**). A P10 Δ3a7b-Nluc passage in the presence of PBS, without nirmatrelvir, was sequenced as an internal control (P10 PBS) to test for cell-culture adaptations, as well as the P0 Δ3a7b-Nluc virus. Amino acid changes with a variant frequency exceeding 20% are presented in **Fig. 5A**. No mutations were identified in the P0 Δ3a7b-Nluc virus, while five mutations were identified in the P10 PBS lineage within NSP3 (E1270D), NSP4 (A260V and M458I), NSP15 (G13E), and E (R38Q). In contrast, eight mutations, including NSP2 (K67E), NSP5 (L50F and R188G), NSP6 (A136V and L260F), NSP10 (D82G), NSP13 (V6F), and E (S68F) were identified in P10 Δ3a7b-Nluc DRM-N. Because nirmatrelvir targets Mpro, mutations L50F and R188G in NSP5 gene were considered leading candidates for causing the drug-resistance phenotype. To validate the identified mutations in NSP5 gene, RNA samples from Vero-AT cells infected with P10 Δ3a7b-Nluc DRM-N were subjected to RT-PCR and Sanger sequencing (**Figs. 5B-5C**). Sequencing results were consistent with NGS data, confirming the presence of L50F (**Fig. 5B**) and R188G (**Fig. 5C**) in NSP5 gene.

**Figure 5.**
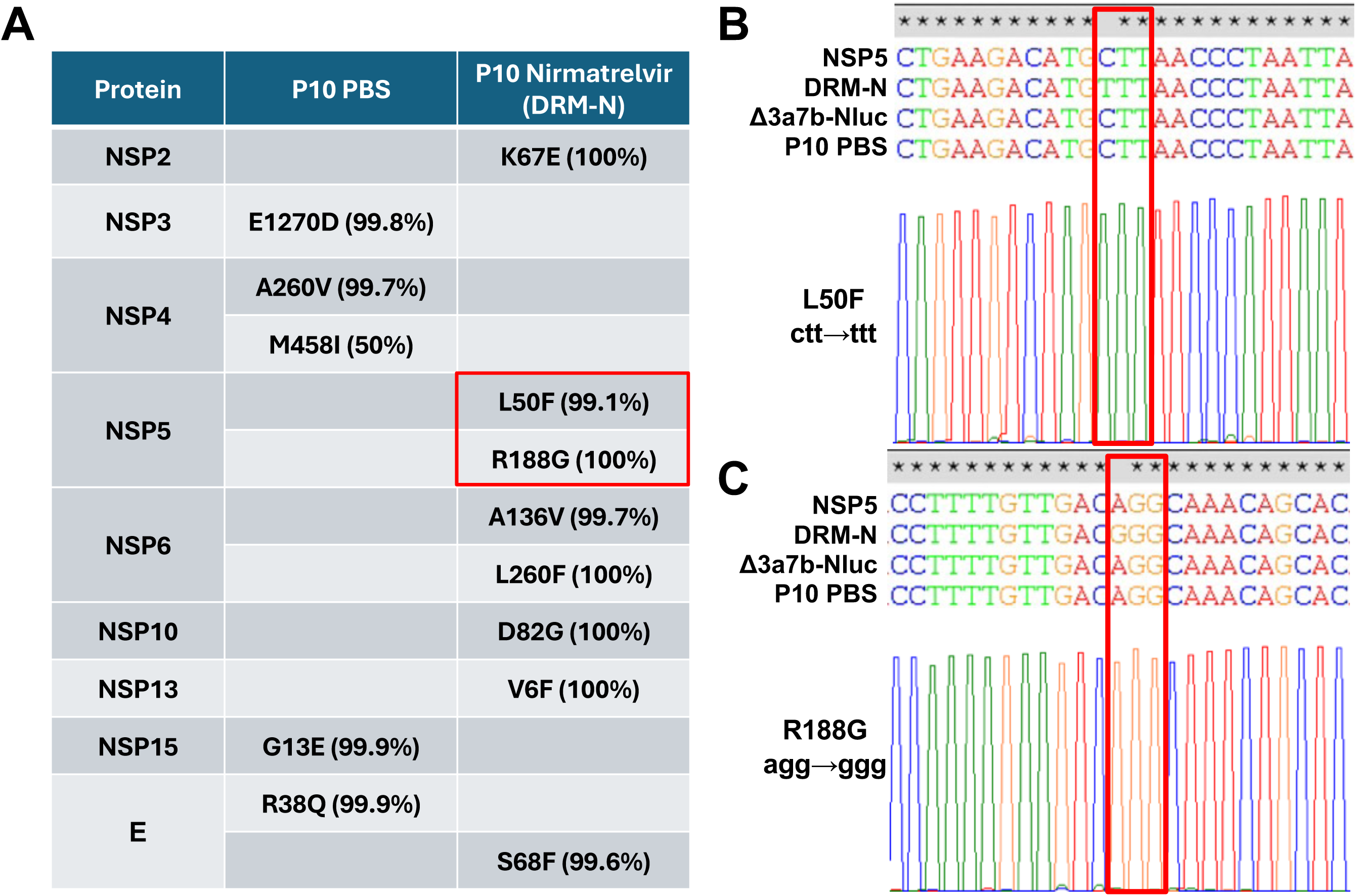
Identification of P10 Δ3a7b-Nluc DRM-N mutations. (A) NGS was performed on RNA isolated from Δ3a7b-Nluc that was cultured in the presence of PBS or increasing concentrations of nirmatrelvir for 10 serial passages, P10 PBS and P10 DRM-N, respectively. Identified amino acid changes and their protein locations are indicated. DRM-N mutations L50F and R188G located in NSP5 gene are indicated with a red rectangle. (B-C) RNA used for NGS was used for RT-PCR amplification of NSP5 as well as for Sanger sequencing to confirm the presence of L50F (B) and R188G (C) mutations in P10 Δ3a7b-Nluc DRM-N, which are indicated with red rectangles.

### Characterization of mutations identified in P10 Δ3a7b-Nluc DRM-N

To investigate the contribution of the identified mutations to nirmatrelvir resistance, we generated Δ3a7b-Nluc variants containing L50F, R188G, and L50F+R188G mutants. The plaque morphology of Δ3a7b-Nluc L50F, R188G and L50F+R188G was compared to that of Δ3a7b-Nluc in Vero-AT cells (**Fig. 6A**). The Δ3a7b-Nluc R188G mutant exhibited smaller plaque diameters compared to Δ3a7b-Nluc, whereas the Δ3a7b-Nluc L50F mutant developed slightly larger plaques than Δ3a7b-Nluc. Notably, the L50F+R188G mutant showed a plaque size resembling that of Δ3a7b-Nluc, suggesting that L50F may compensate the smaller plaque phenotype observed with Δ3a7b-Nluc R188G (**Fig. 6A**). The growth kinetics of the different mutants were next compared in Vero-AT cells infected at 0.01 MOI. Relative to Δ3a7b-Nluc, R188G exhibited a growth delay at 24-, 48- and 72-hours post-infection and the L50F+R188G showed an intermediate growth phenotype (**Fig. 6B**). We next evaluated the contribution of each mutant to nirmatrelvir resistance by determining EC_50_ values of the mutant and original Δ3a7b-Nluc viruses using the Nluc assay. Both L50F and R188G viruses showed ∼5-fold increases in nirmatrelvir resistance (EC_50_ = 0.56 µM for Δ3a7b-Nluc, 2.80 µM for L50F, and 2.69 µM for R188G) (**Fig. 6C**). Combination of L50F and R188G resulted in an EC_50_ of 6.07 µM, representing an approximately 11-fold increase from the parental Δ3a7b-Nluc (**Fig. 6C**). Importantly, all viruses showed similar EC_50_ values to remdesivir, confirming that the increased drug tolerance was specific to nirmatrelvir (**Fig. 6D**).

**Figure 6.**
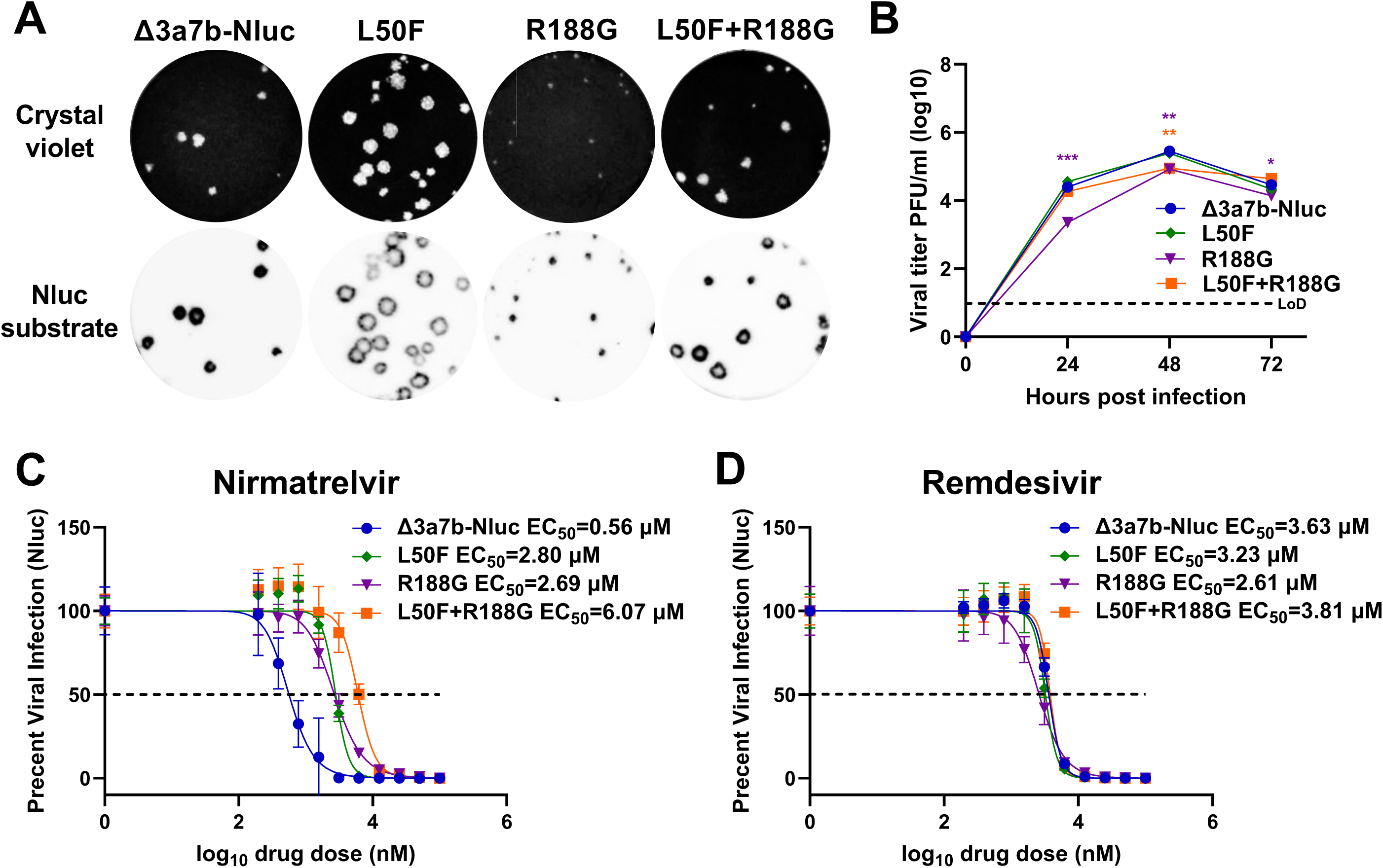
Characterization of Δ3a7b-Nluc L50F, R188G, and L50F+R188G mutants. Vero-AT cells (6-well plate format) were infected with Δ3a7b-Nluc parental, L50F, R188G, and L50F+R188 viruses. At 72 hours post-infection, plates were incubated with crystal violet (top) or stained with Nluc substrate (bottom). (B) Vero-AT cells (6-well plate format, triplicates) were infected (MOI 0.01) with Δ3a7b-Nluc parental, L50F, R188G and L50F+R188 viruses. At the indicated hours post-infection, cell culture supernatants were collected, and presence of virus was determined by plaque assay. LoD : limit of detection. The antiviral activity of nirmatrelvir (C) and remdesivir (D) against parental Δ3a7b-Nluc and DRM-N L50F, R188G and L50F+R188 viruses was determined in Vero-AT cells (96-well plate format, quadruplicates) by Nluc activity. EC_50_ values of the different antivirals were calculated using GraphPad Prism. Data are presented as means ± SD. Dotted line indicates the 50% of viral inhibition. Data are presented as mean ± SD. The statistical significance was analyzed with one-way ANOVA followed by Tukey’s test (* P < 0.05, ** P < 0.01, *** P < 0.001).

## DISCUSSION

Despite the development of vaccines, monoclonal antibodies, and antiviral drugs, the emergence of SARS-CoV-2 variants continues to threaten public health globally (6–13). Mutations in the Spike (S) protein have challenged the efficacy of the original vaccines and undermined antibody therapeutics (6–13). Mutations in NSP5 gene could potentially become a challenge to protease inhibitors such as paxlovid and ensitrelvir, since laboratory-derived and naturally occurring antivirals-escape mutations have been identified (28–33). Further characterizing the DRM landscape may guide the development of effective antiviral strategies against SARS-CoV-2.

We generated an attenuated rSARS-CoV-2 lacking ORFs 3a and 7b and expressing Nluc (Δ3a7b-Nluc) to establish a safe platform to isolate and characterize drug-resistant mutants without the biosafety concerns of conducting adaptation experiments with WT forms of SARS-CoV-2. Deletion of ORFs 3a and 7b provides a bona-fide surrogate to investigate drug resistance while expression of Nluc allows easy tracking of viral infection. Moreover, the use of Δ3a7b-Nluc provides a safe approach to verify candidate resistance mutations. Thus, Δ3a7b-Nluc can be used to easily and quickly identify and validate drug-resistant variants.

In this study, we identified two specific mutations, L50F and R188G, within the Mpro of SARS-CoV-2 responsible for resistance to nirmatrelvir. Neither mutation is directly involved in the binding of nirmatrelvir to Mpro (**Fig. 7**). Instead, both amino acid residues located in the hydrophobic S2 pocket. L50F has been previously reported to cooperate with an E166V mutation to confer resistance to nirmatrelvir (17, 28, 29). The substitution of leucine (L) to phenylalanine (F) introduces a bulky side chain, strengthening hydrophobic interactions between F50 and Q189 and stabilizing the main chain of Q189. Additionally, the L50F substitution made the S2 pocket more compact, which could facilitate hydrophobic interactions between Mpro and its substrate, resulting in increased enzymatic activity of Mpro. The R188G mutation found in our study is also located in the S2 hydrophobic pocket (**Fig. 7**). This substitution replaces the highly hydrophilic arginine (R) with a less hydrophilic glycine (G), which is also predicted to increase the compactness of the S2 pocket. Notably, the R188G mutation impairs viral proliferation, while the L50F mutation seems to compensate for this loss of activity. A similar compensatory mechanism was observed with other mutation combinations, T21I+S144A and T21I+E166V (28). It was found that T21I primarily contributes to fitness compensation, whereas S144A or E166V directly interfere with the binding of nirmatrelvir to Mpro (17). These findings demonstrate that although SARS-CoV-2 can develop resistance to nirmatrelvir through multiple mechanisms, the available resistance landscape appears to be limited. Previous studies also identified the R188G mutation, but not in combination with L50F (28). Reportedly, a rSARS-CoV-2 L50F was 4.2-fold less sensitive to nirmatrelvir than WT SARS-CoV-2 (28), which is similar to the ∼5-fold change that we observed in this study with Δ3a7b-Nluc.

**Figure 7.**
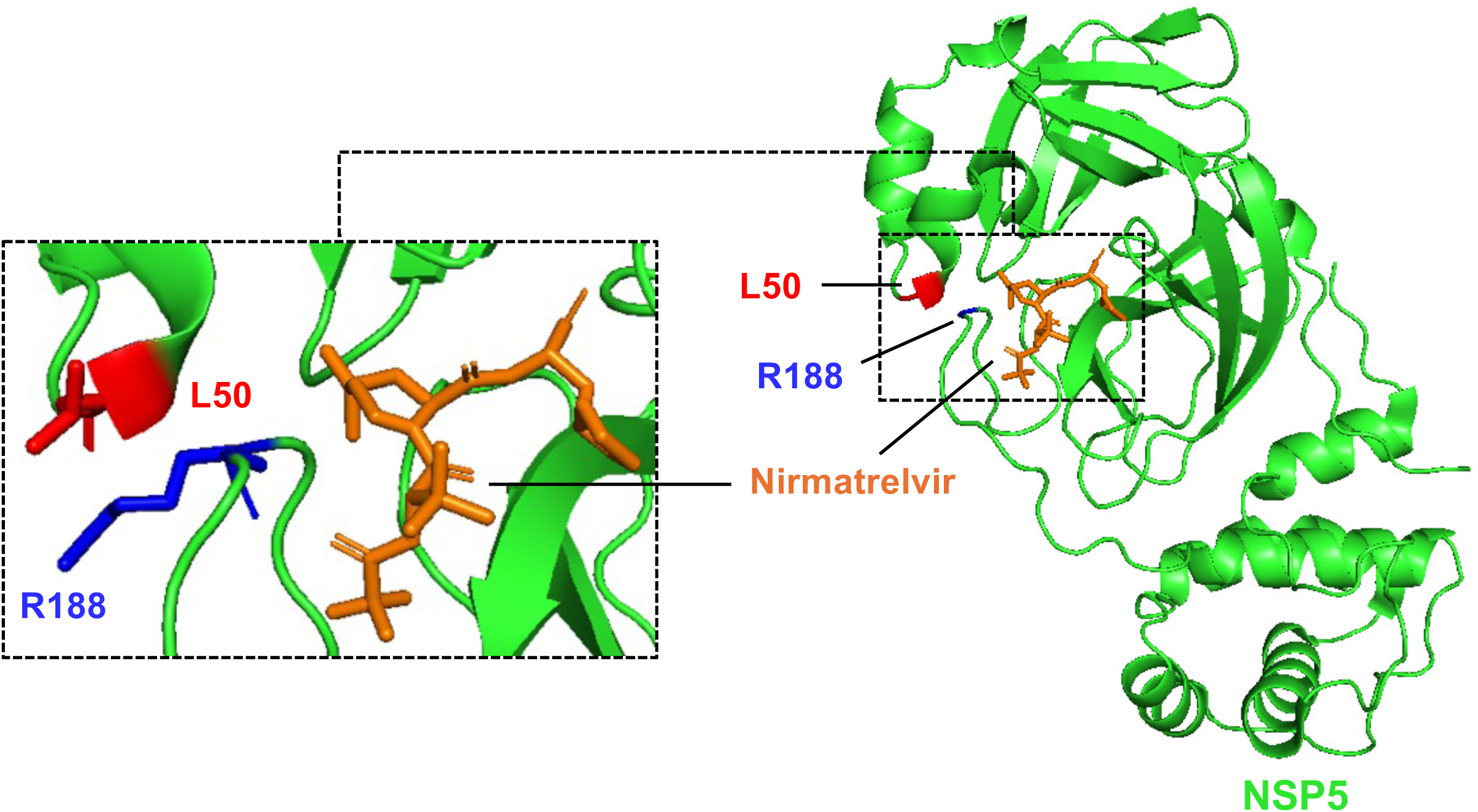
Schematic ribbon diagram of Mpro-nirmatrelvir binding. Amino acids L50 (red) and R188 (blue) in Mpro (green) are shown. Nirmatrelvir is indicated in orange. The Mpro-nirmatrelvir complex was obtained from PDBID: 7VH8.

In conclusion, our study demonstrates the feasibility of using the attenuated Δ3a7b-Nluc as a safe and effective approach for identifying and validating SARS-CoV-2 drug-resistance mutations to antivirals without the biosafety concern of conducting adaptation experiments with WT SARS-CoV-2. The identification of nirmatrelvir L50F and R188G resistance mutations provides valuable insights into the mechanism of viral escape from paxlovid and highlights the need for ongoing development of additional therapeutics. In addition to resistance profiling, Δ3a7b-Nluc could be used to reveal viruses with resistance to immune responses induced by vaccination without GoF concerns. Thus, Δ3a7b-Nluc may aid the rapid development of next-generation therapeutics or prophylactics to improve preparedness against further SARS-CoV-2 evolution.

## MATERIALS AND METHODS

### Biosafety

All experiments utilizing SARS-CoV-2 were conducted in Biosafety Level 3 (BSL-3) containment laboratories at the Texas Biomedical Research Institute (Texas Biomed) and approved by the Institutional Biosafety Committee (IBC). Experiments with Δ3a7b and Δ3a7b-Nluc were conducted at BSL-2+.

### Cells

Vero cells expressing hACE2 and TMPRSS2 (Vero-AT) were obtained from BEI Resource and maintained in Dulbecco’s modified Eagle medium (DMEM) supplemented with 10% fetal bovine serum (FBS; VWR), 100 units/ml penicillin-streptomycin (Corning) and puromycin 10 µg/ml (InvivoGen).

### Recombinant viruses

The rSARS-CoV-2 Δ3a7b-Nluc (Δ3a7b-Nluc) was rescued in Vero-AT according to our previously described protocol (35, 36). Briefly, the BAC containing the Δ3a7b-Nluc genome was verified by restriction enzyme digestion and used to transfect confluent Vero-AT cells (6-well plates) using Lipofectamine 2000 (Thermo Fisher Scientific). After 24 hours, the medium was replaced with post-infection medium. At 48 hours post-transfection, cells were scale up into T75 flasks. After incubation for another 72 hours, cell culture supernatants were collected and stored at -80°C. The rSARS-CoV-2 WT and the rSARS-CoV-2 lacking ORFs 3a and 7b (Δ3a7b) were previously described (34).

### Plaque assay and immunostaining

Monolayer of Vero-AT cells (6-well plate) were infected with 10-fold serial dilutions of the indicated viruses for 1 hour at 37°C in a 5% CO_2_ incubator. After viral adsorption, cells were overlaid with media containing 1% agar and incubated at 37°C in a 5% CO_2_ incubator for 72 hours. Then, cells were fixed overnight with 10% formaldehyde solution. For Nluc visualization, plates were treated with Nano-Glo Luciferase Assay substrate (Promega) and imaged in a ChemiDoc MP Imaging System. For immunostaining, cells were permeabilized with 0.5% Triton X-100 in PBS for 15 minutes at room temperature (RT), followed by staining with the SARS-CoV cross-reactive N protein 1C7C7 monoclonal antibody (1 μg/mL), the Vectastain ABC-HRP kit (Vector Laboratories), and DAB Substrate Kit (Vector Laboratories). Plates were scanned and photographed using a ChemiDoc MP Imaging System. Finally, cells were stained with crystal violet and imaged with the ChemiDoc MP Imaging System.

### Growth kinetics

Confluent Vero-AT cells (6-well plate, triplicates) were infected (MOI 0.01) with viruses at 37°C in a 5% CO_2_ incubator for 1 hour. After viral adsorption, cells were washed with PBS and incubated with post-infection medium (DMEM with 2% FBS, 100 units/ml penicillin-streptomycin) at 37°C in a 5% CO_2_ incubator. At the indicated times post-infection (24, 48, and 72 hours), viral titers were determined by plaque assay and immunostaining as described above. Presence of Nluc in the cell culture supernatant was quantified using Nano-Glo Luciferase Assay System.

### Isolation of DRM-N

Confluent monolayers of Vero-AT cells (12-well plates, triplicates) were infected with Δ3a7b-Nluc (100∼200 PFU/well) for 1 hour at 37°C in a 5% CO_2_ incubator. After viral adsorption, cells were washed with PBS and cultured in the presence of nirmatrelvir (starting concentration of 1 µM). After 72 hours, the cell culture supernatant of the highest nirmatrelvir concentration with more than 50% cytopathic effect (CPE) was collected and titrated by Nluc activity for next passage. After 10 serial passages in increasing concentration of nirmatrelvir, P10 Δ3a7b-Nluc DRM-N was collected, amplified in Vero-AT cells and aliquoted at -80°C.

### Immunofluorescence assay

Vero-AT cells (6-well plates) were infected (MOI 0.001) with the indicated viruses at 37°C in a 5% CO_2_ incubator for 1 hour. After viral adsorption, virus inoculum was replaced with post-infection medium containing the indicated concentrations of nirmatrelvir (0 µM, 2.5 µM, 10 µM, and 40 µM) and cultured at 37°C in a 5% CO_2_ incubator. After 48 hours infection, cells were fixed and permeabilize, and presence of virus was determined using the cross-reactive SARS-CoV N protein antibody 1C7C7 and an anti-mouse FITC-secondary antibody. The cell nucleus was stained with DAPI. Cells were visualized and imaged under an EVOS fluorescent microscope (Thermo Fisher Scientific).

### Half-maximal inhibitory concentration assay

Plaque reduction neutralization test (PRNT): Confluent Vero-AT cells (96-well plates, quadruplicates) were infected with 100∼200 PFU/well of Δ3a7b-Nluc and incubated at 37°C in a 5% CO_2_ incubator for 1 hour. The virus inoculum was replaced after viral adsorption and cells were incubated in post-infection medium containing serial dilutions of nirmatrelvir (starting concentration of 100 µM) and 1% Avicel, followed by incubation at 37°C in a 5% CO_2_ incubator. At 18 hours post-infection, cells were fixed with 10% formalin for 24 hours. After washing with PBS, cells were permeabilized with 0.5% Triton X-100 at RT for 15 minutes. After washing, cells were stained with the SARS-CoV N protein 1C7C7 monoclonal antibody, the Vectastain ABC-HRP kit, and the DAB Substrate Kit according to the manufacturer’s instructions. Plates were scanned and imaged using a ChemiDoc MP Imaging System.

Nluc assay: Confluent Vero-AT cells (96-well plates, quadruplicates) were infected with 100∼200 PFU/well of Δ3a7b-Nluc and incubated at 37°C in a 5% CO_2_ incubator. After 1 hour adsorption, virus inoculum was replaced with post-infection medium containing serial dilutions of nirmatrelvir (starting concentration of 100 µM) and cultured at 37°C for 48 hours. Cell culture supernatants were then collected and assessed for the presence of Nluc using the Nano-Glo Luciferase Assay substrate. Nluc was measured using a GloMax Discover system (Promega).

Crystal violet staining assay: Confluent Vero-AT cells (96-well plates, quadruplicates) were infected with 100∼200 PFU/well of Δ3a7b-Nluc and incubated at 37°C in a 5% CO_2_ incubator. After 1 hour adsorption, the virus inoculum was replaced with post-infection medium containing serial dilution of nirmatrelvir (starting concentration of 100 µM) and incubated at 37°C in a 5% CO_2_ incubator for 72 hours. Then, cell culture supernatants were removed, and the plates were fixed with 10% formalin solution for 24 hours. After washing with PBS, cells were stained with crystal violet ON. Plates were rinsed with water for 5 times, and 100 µl of 100% methanol was added. After 20 minutes incubation at RT, the optical density at 560 nm was measured using a GloMax Discover system.

### Sequencing

Total RNA was extracted from infected Vero-AT cells using Trizol reagent (Thermo Fisher Scientific) following the manufacturer’s recommendations. Metagenomic NGS was conducted as described previously (39). Briefly, RNA was treated with Turbo DNAse, reverse transcribed using random hexamers using SuperScript IV (Thermo Fisher), and double-stranded cDNA was created using Sequenase v2.0 (Thermo Fisher). Double-stranded cDNA was tagmented using Illumina DNA Prep (S) kit and 14 cycles of dual-indexed PCR and sequenced using a 2x150bp run on a NovaSeq 6000. Reads were trimmed and quality-filtered using fastp (40). Variant analysis was performed using RAVA (using the rWA1 Δ3a7b-Nluc reference sequence. Briefly, reads were aligned with bwa-mem (41), variants called with samtools and annovar (42, 43), and results presented using custom visualization. RNA extracted from infected Vero-AT cells was also reverse-transcribed into cDNA using SuperScript II reverse transcriptase (Thermo Fisher Scientific). The NSP5 gene was amplified by PCR using the Expand high-fidelity PCR system (Sigma-Aldrich), and the resulting products were Sanger sequenced by Plasmidsaurus. Primer sequences are available upon request.

### Statistical analysis

All data represent the means ± standard deviation (SD) for each group and were analyzed with GraphPad Prism. The statistical significance was analyzed with one-way ANOVA followed by Tukey’s test. P values <0.05 were regarded as statistically significant.

## Data availability

Sequencing reads for viral whole genome sequencing are available in NCBI BioProject PRJNA1255607.

## ACKNOWLEDGMENTS

We want to thank BEI Resources for providing Vero-AT cells. This work was supported, in part, by public health service grant U19 AI171403 from the NIH/NIAID (to RKP). All experimental work in this study supported by this award was completed at or before March 24, 2025.

